# Genomic comparison of highly related pairs of *E. coli* and *K. pneumoniae* isolated from faeces and blood of the same neonatal patients hospitalized with fever in Dar es Salaam, Tanzania

**DOI:** 10.1101/2025.04.28.650962

**Authors:** Richard N. Goodman, Sabrina J. Moyo, Ilinca Memelis, Aakash Khanijau, Joel Manyahi, Upendo O. Kibwana, Said Aboud, Bjørn Blomberg, Nina Langeland, Adam P. Roberts

## Abstract

Blood stream infections (BSIs) are a major cause of hospitalisation and death for children under the age of five in sub-Saharan Africa with members of the Gram-negative bacteria Enterobacterales such as *Klebsiella pneumoniae* and *Escherichia coli* among the most common causative agents. These bacteria usually colonise the human gastrointestinal (GI) tract which has been identified as a reservoir for invasive infections into extra-intestinal environments such as the urinary tract and bloodstream. In this study we used comparative genomics to compare hybrid genome assemblies of blood and faecal isolates taken from the same patients (all neonates under 19 days old) to determine if the BSI associated bacterial isolates originated in their GI tract. We show that both *E. coli* and *K. pneumoniae* likely translocated from the GI tract to the blood in multiple cases of BSI. We also highlight key virulence genes and acquired mutations that are indicative of pathogenic strains capable of BSI. These findings expand our understanding of BSI pathogenesis and could help guide targeted interventions to prevent future BSI infections in neonates.

## Introduction

Blood stream infections (BSI) are a major cause of sepsis, hospitalisation and death for children under the age of 5. Worldwide there was an estimated 2.9 million sepsis related deaths in 2017 with the highest burden in sub-Saharan Africa ^1^. In this region neonates can be particularly susceptible to sepsis caused by invasive bacterial infections due to co-morbidities such as HIV infection, malnutrition, malaria and sickle-cell disease ^2–4^. It is often difficult to distinguish between febrile illness in infants caused by bacteria and malaria or other infectious agents within low-middle-income countries (LMICs), making them hard to diagnose and treat ^5–7^. Gram-negative bacteria, including the Enterobacterales *Klebsiella pneumoniae* and *Escherichia coli*, were shown to be the main causative agents of neonatal sepsis in a study of several LMICs in Africa and South Asia ^8^.

*K. pneumoniae* and *E. coli* both make up part of the human, commensal, intestinal microbiota, and colonisation of the neonatal gastrointestinal (GI) tract with such organisms occurs rapidly after birth, influenced by the maternal flora and peri-partum environment. In the context of neonatal sepsis, the colonised GI tract has been identified as a reservoir for BSI ^9^. This can arise from direct bacterial translocation from the gastrointestinal (GI) tract to the blood across the gut epithelium, or via secondary colonisation and infection of other body sites such as the skin, respiratory or urinary tract ^10,11^. Bacterial translocation through the intestinal mucosal barrier into the bloodstream can occur during gut dysbiosis ^12^ or increased intestinal permeability ^13^. Premature neonates and those born with low birth weight are particularly vulnerable, as immature immune and gastrointestinal systems can lead to breach of the mucosal barrier ^11,14,15^, and antibiotic exposure seen in healthcare environments such as the neonatal intensive care unit can contribute to gut dysbiosis^16^. Once the intestinal mucosal barrier has been traversed and bacteria are in the blood, this can lead to sepsis. To survive within two distinct environments as diverse as the intestine and blood, *K. pneumoniae* and *E. coli* isolates may already contain all the genes they need, or they may require adaptive mutations or the acquisition of genes via mobile genetic elements^17^.

A previous study investigated the trends of bacteraemia amongst 2,226 children under 5 years of age who were hospitalised with fever in Dar es Salaam, Tanzania ^7^. This study builds on that work, by directly comparing bacterial isolates taken from blood and faecal samples of individual neonates on the same day, to assess whether the faecal and blood isolates are likely related. We started with the hypothesis that *E. coli* and *K. pneumoniae* inhabiting the gut microbiome translocated to the blood before the onset of the BSI and associated symptoms.

We used comparative genomics to determine differences between the paired blood and faecal isolates taken from the same patients. This enabled us to determine if they were highly related and if there were changes within the genome when the bacterial isolates translocated from the gut to the blood. Using a variety of bioinformatic methods, we show there are both *E. coli* and *K. pneumoniae* that likely translocated to the blood stream and highlight key virulence genes and acquired mutations that may be associated with this translocation.

Studies from the USA and Taiwan have analysed faecal metagenomes of pre-term and very low birth weight neonates showing bacterial translocation in patients ^11,12,18^. There have also been studies which used comparative genomics to compare *E. coli* and *K. pneumoniae* from the urinary tract and gut in patients in the USA ^19,20^. There have been further studies exploring the virulence factors which contribute to bacterial translocation to extra-intestinal sites and subsequent infections in the urinary tract of patients in France ^21^. However, to our knowledge, this is the first study that has directly compared hybrid assembled genomes of blood and faecal *E. coli* and *K. pneumoniae* isolates from the same patients at the genomic and molecular level and highlighted virulence genes and acquired mutations that may contribute to the gut-blood transition within individual patient isolates. This is also the first study to investigate bacterial translocation to extra-intestinal sites in a sub-Saharan Africa setting. The hybrid assemblies provide unique structural and contextual information about genes and replicons allowing an in-depth comparative genomics approach. These findings expand our understanding of BSI pathogenesis from Gram-negative organisms and the dynamic genetic landscape likely involved in the translocation from the gut to the blood.

## Results

### Typing of *E. coli* and *K. pneumoniae* isolated from blood and faeces

In a previous study 2,226 blood-cultures were analysed from children below the age of 5 years hospitalised with fever in Dar es Salaam, Tanzania with 82% of participants under the age of two ^7^. Sixty-two *K. pneumoniae* and 28 *E. coli* were isolated from the blood cultures of these patients, totalling 90 blood isolates. Thirteen of these blood isolates had a paired faecal isolate taken from the same patient ^22^. These blood and faecal isolates were sequenced and hybrid assembled (see Supplementary Table 1) to determine relatedness and determine the likelihood of bacterial translocation from the gastrointestinal (GI) tract to the blood stream (see Table 1). Therefore, in total 26 isolates were assessed as 13 faecal (FC) and blood (BL) pairs; 4 pairs were confirmed as *Escherichia coli* (*n*=8), 8 pairs of *Klebsiella pneumoniae* (*n*=16), and 1 pair of consisting of *Klebsiella pneumoniae* and *Klebsiella quasipneumoniae*^23^ (n=2). These were from patients under the age of 19 days with fevers lasting 1 – 3 days (Supplementary Table 2). When referencing the genomic data, the prefix FS refers to patient ID (e.g. FS2155), the prefix FSBL refers to the blood isolate (FSBL2155) and the prefix FSFC refers to the faecal isolate (FSFC2155).

**Table 1:**
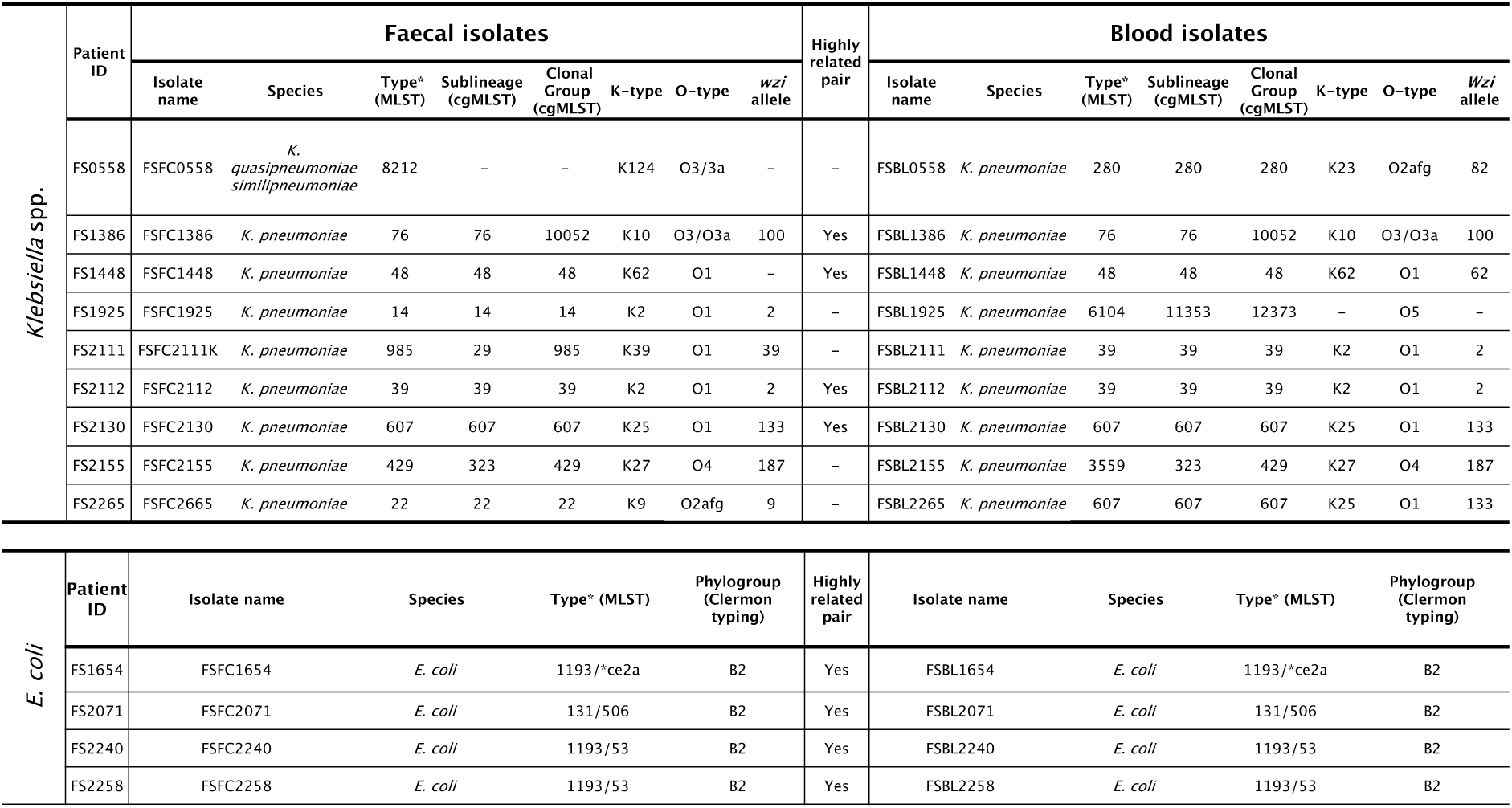
MLST profiles of the 13 paired blood and faecal isolates (n = 26). Multi-locus sequence typing (MLST) from Pasteur Institute, France for *Klebsiella* species (https://bigsdb.pasteur.fr/klebsiella/) and Warwick University, UK for *E. coli* (http://mlst.warwick.ac.uk/mlst/dbs/Ecoli). Core genome MLST (cgMLST) from Pasteur Institute, France for *Klebsiella* species (https://bigsdb.pasteur.fr/klebsiella/cgmlst-lincodes/). O-types, K-types and *wzi* allele for *Klebsiella* species were predicted with Kaptive ^24^. For *E. coli* phylogroups were assigned using ClermonTyping ^25^. The highly related pairs indicate shared MLST and, for *Klebsiella* spp., cgMLST. FS2155 shared the same cgMLST but not MLST so is not defined as a highly related pair.

Firstly, the relatedness of the paired blood and faecal isolates was assessed using multi-locus sequence typing (MLST) and core genome MLST (cgMLST). Eight out of the thirteen pairs (4 *E. coli* and 4 *K. pneumoniae*) shared the same MLST and nine shared the same cgMLST (4 *E. coli* and 5 *K. pneumoniae*). The discrepancy occurred with patient FS2155, the faecal (FSFC2155) and blood (FSBL2155) MLST profiles were ST426 and ST3559, respectively, but the sublineage (323) and clonal group (ST429) from cgMLST were the same for both as well as the K-type, O-type and *wzi* capsular gene type (see Table 1). Of the four *K. pneumoniae* highly related pairs, all had the same within-pair K-type, O-type and *wzi* gene type. Of the four *E. coli* pairs all isolates, both from blood and faeces were of the phylogroup B2. The *E. coli* isolates from patient FS2071 belonged to ST131 and the *E. coli* isolates from patients FS1654, FS2246 and FS2258 belonged to ST1193.

### Determining genome similarity between paired isolates using Average Nucleotide identity (ANI) and core genome single nucleotide polymorphisms (SNPs)

To further assess the relatedness of the paired blood and faecal, average nucleotide identities (ANI) and core genome SNP (SNPs) distances were determined for *E. coli* and *Klebsiella* spp. (Figure 1) and between all isolates (Supplementary Figure 1). Of the 9 pairs sharing the same MLST, 8 had an ANI of 99.99% or above and < 23 SNPs, with FS2155 isolates having an ANI of 99.78 and 7 SNPs (see Figure 1). The ANI analysis also revealed close relatedness between isolates which were not part of the same pair. For example the FSBL2111 isolate shared an ANI of 100% and 0 SNPs with both FSFC2112 and FSBL2112 indicating they are all the same strain. FSBL2265 shares an ANI of 99.88% with FSBL2130 (0 SNPs) and 100% (0 SNPs) with FSFC2130 indicating they are the likely the same sequence type or possibly the same strain. FSBL1654 and FSBL2240 share an ANI of 100% with FSFC2258 (see Figure 1).

**Figure 1:**
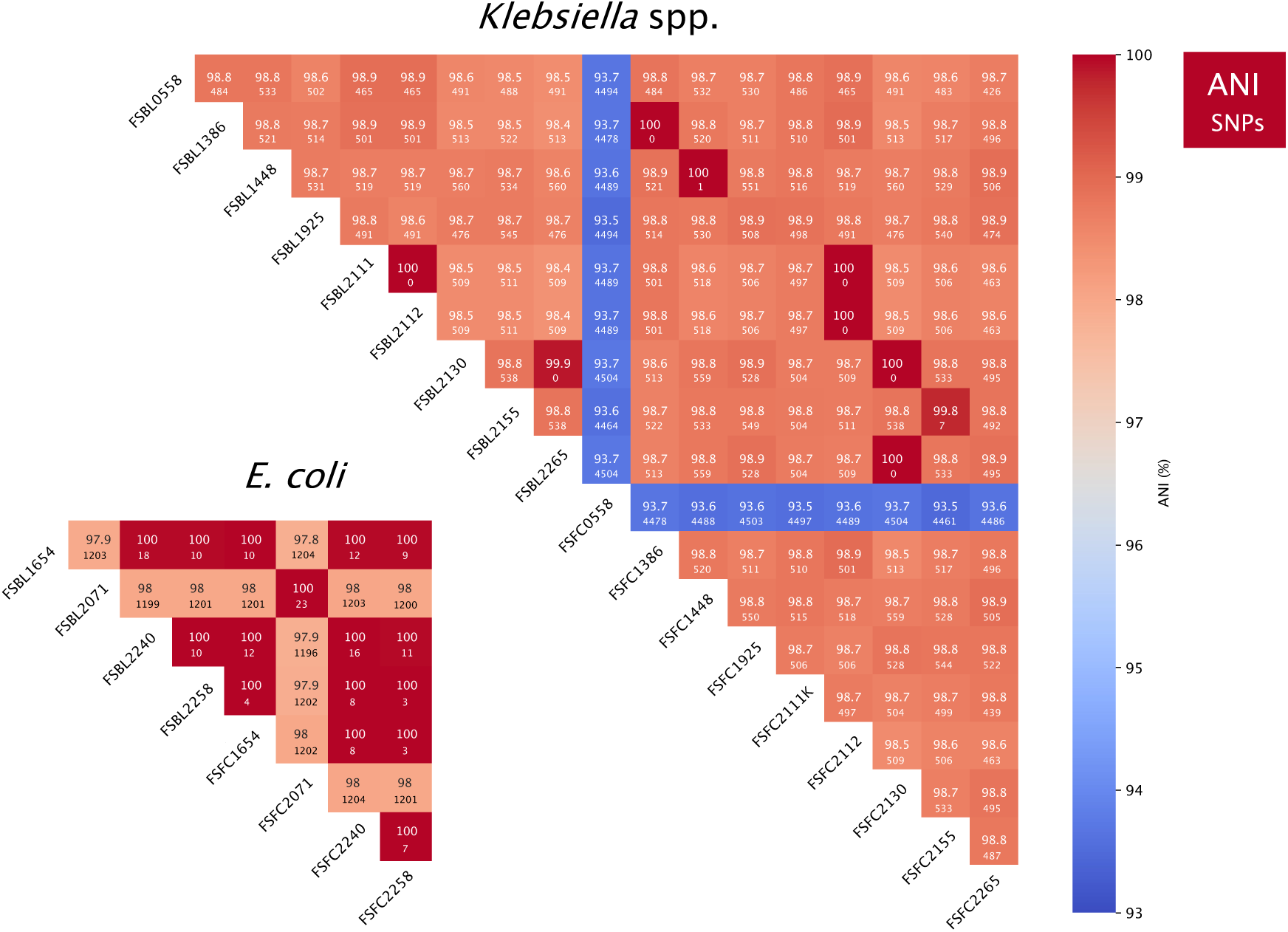
Average nucleotide identities (ANI) and core genome single nucleotide polymorphisms (SNPs) across 4 *E. coli* and 9 *Klebsiella* spp. paired isolates. ANI was calculated using the BLAST alignment algorithm (ANIb). The core genome SNPs are displayed in each cell below the ANI %, this was calculated using snippy v4.6.3 and snp-dist using FSFC1386 as reference (see methods). *Klebsiella* spp. are displayed together with 17 *Klebsiella pneumoniae* and 1 (FSFC0558) *Klebsiella quasipneumoniae* highlighted by the lines of blue.

### Phylogenetics reveals the genomic relatedness across isolates

Core genome trees were built based on SNP differences between the core genomes of closely related isolates (see Figure 2A), all 8 pairs that share the same MLST and share an ANI above 99.99% were on the same branch with distances less than 0.00008 branch length between nodes. The isolates from FS2155 were also on the same branch. As with the ANI analysis, isolates from different patients were closely related, this included FSBL2111 which was on the same branch as FSBL2112 and FSFC2112, and FSBL2265 which was on the same branch as FSFC2130 and FSBL2130. Further to this, both blood and faecal isolates from FS1654, FS2240 and FS2258 were closely related on the tree.

**Figure 2:**
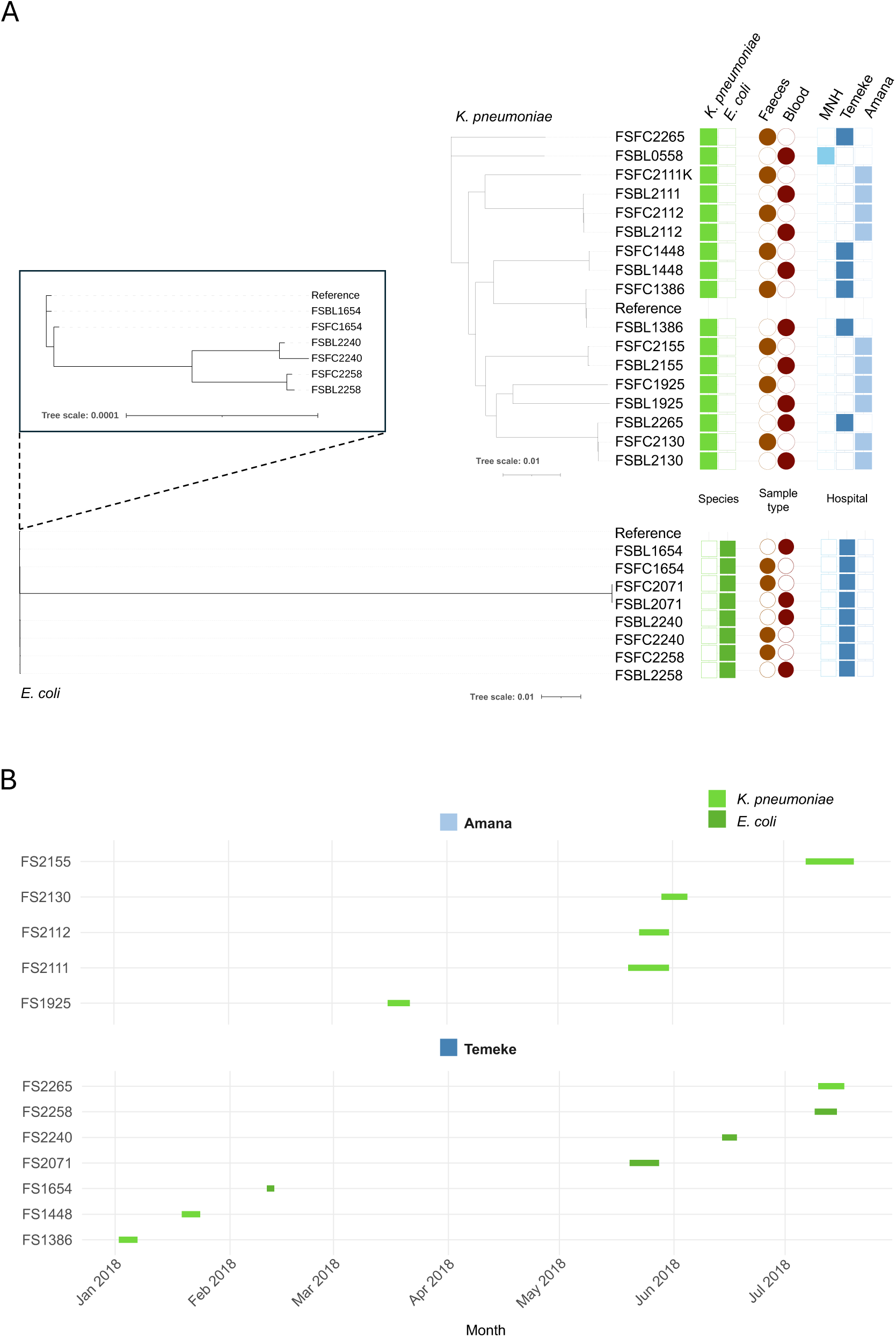
Core genome phylogenetic trees and admission timelines reveal relatedness across isolates. A) Core genome phylogenetic trees of *K. pneumonia*e and *E. coli* isolates. The phylogenetic trees are based on SNPs derived from the core genome alignment of *K. pneumonia*e and *E. coli* isolates. The *Klebsiella pneumoniae* subsp. *pneumoniae* HS11286 genome (GCF_000240185.1) was used as reference for *Klebsiella pneumoniae* isolates and the *Escherichia coli* str. K-12 substr. MG1655 genome (GCF_000005845.2) was used as reference for *E. coli* isolates. Branch lengths are representative of the numbers of SNPs between isolates. For the *E. coli* tree FS2071 was removed to clearly visualise the relationship between the other paired isolates (shown in black box). The coloured shapes indicate metadata such as species (green), sample type (brown/red) and hospital of origin (blue). MNH; Muhimbili National Hospital. B) Timelines from admission to discharge of neonatal patients in Temeke and Amana hospitals are represented by the start and end of the green lines in the timeline. MNH was excluded as only one patient was from this study site. The colours represent species consistent with the panel above.

Isolates from patients FS2111 and FS2112 were taken from the same hospital (Amana) within 3 days of each other (see Figure 2B) indicating possible nosocomial transmission. Isolates from patient FS2265 came from Temeke hospital and isolates from patient FS2130 came from Amana, so the similarity in sequence type is likely only incidental. Isolates from patients FS2258, FS1654 and FS2240 were all taken from Temeke hospital but in different months of the year so nosocomial transmission here would indicate an endemic strain within the hospital.

Subsequently, the eight pairs (4 *E. coli* and 4 *K. pneumoniae*) shown to be the same strain from MLST, ANI and SNP-based phylogeny (i.e. isolates from FS1386, FS1448, FS1654, FS2071, FS2112, FS2130, FS2240 and FS2258) will be referred to as highly related pairs. Together, it is likely each of these highly related pairs represent bacterial translocation of the same isolate from the GI tract to the blood stream.

### Comparison of AMR, virulence and metal resistance genes between highly related blood and faecal paired isolates

We then looked to compare the antimicrobial resistance (AMR), virulence factor and metal resistance genes between the highly related pairs to determine whether any genes were acquired or lost in the translocation from gut to blood. The acquired AMR profiles did change in two highly related pairs (see Supplementary Table 3). The percentage identity to the database reference gene changed for *sul2* and *aph(3”)-lb* in FS1448 isolates and there was a gain of *dfrA17* on an integron in FSBL1654 compared to FSFC1654, the *dfrA17* gene confers resistance to trimethoprim (part of co-trimoxazole) (Supplementary figure 2). There were no differences in the genes conferring resistance to amoxycillin, gentamicin and ceftriaxone, which were the main antibiotics used to treat the patients (see Supplementary Table 2). All 9 highly related pairs shared the same virulence factors when compared against the VFDB database and metal resistance genes when compared against the MEGARES database (see Supplementary Table 3).

### Comparative genomics of SNPs between highly related blood and faecal paired isolates

We used breseq to run a genome comparison using the hybrid assembled faecal genome as reference and the long reads of the blood isolates as query for each of the highly related pairs which had been confirmed to be highly related. This identified SNPs (both synonymous and non-synonymous), small INDELs (<100bp) in the coding region and intergenic mutations which had been acquired in the blood isolate compared to the faecal isolate are shown in Figure 3. The number of predicted mutations ranged from 1 to 516 across the nine pairs (Supplementary Table 4), with a median of 10, an average of 77.6 and standard deviation of 167. However, for some isolates large numbers of SNPs were concentrated within a single CDS, for example in isolates from FS2071 there were 97 SNPs between the faecal and blood isolates with 19 in a gene encoding a phage lysozyme, 5 in a gene encoding a phage endopeptidase, and 16 in a gene encoding a phage tail fibre protein.

**Figure 3:**
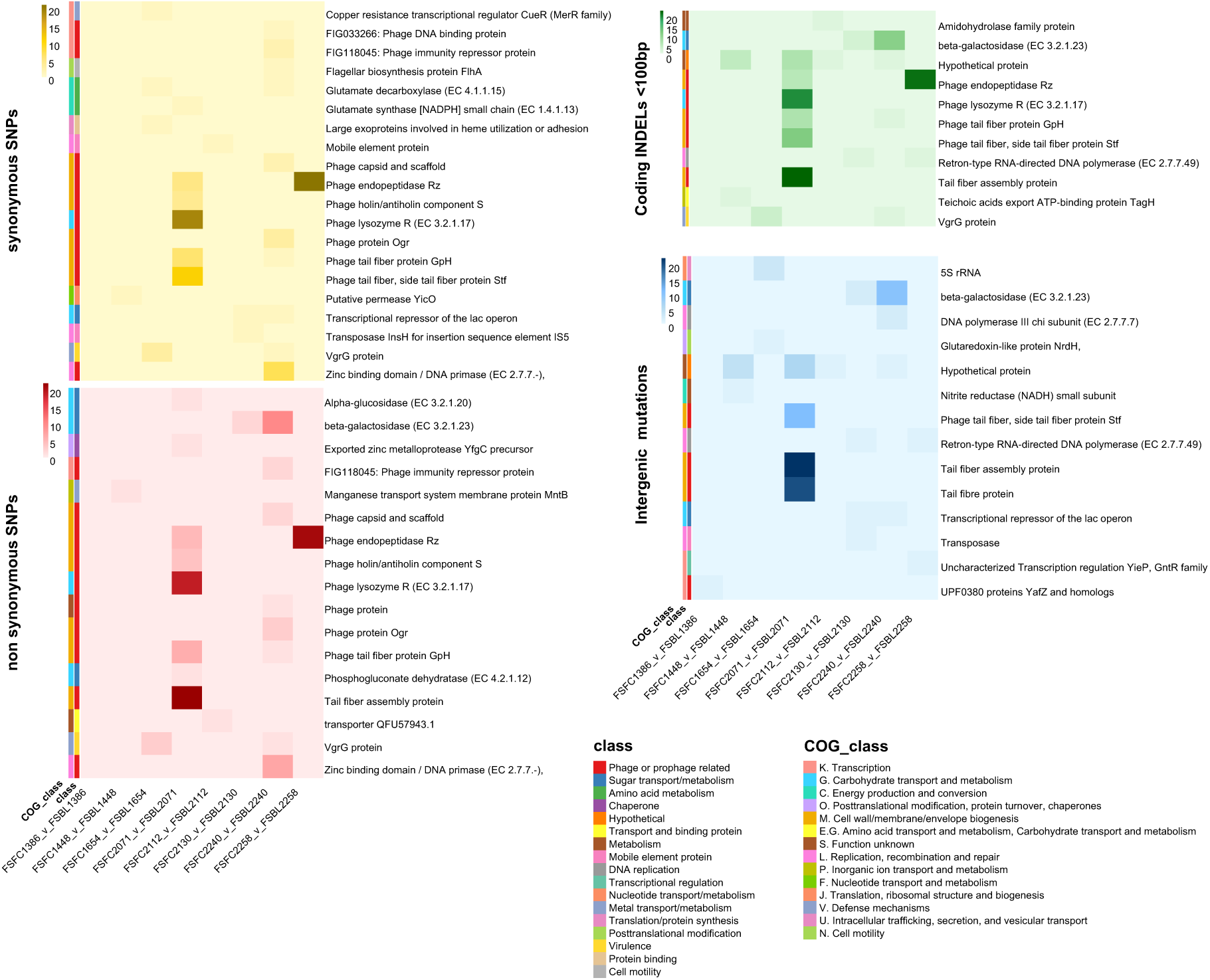
Predicted SNP mutations when comparing the blood genome against the faecal genome for a set of eight highly related paired isolates. All pairs compared with SNP mutations and INDELs <100bp categorised by gene class, gene context and eggNOG clusters of orthologous genes (COG). SNPs were identified with breseq. Heatmaps were visualised separately for synonymous SNPs, non-synonymous SNPs, INDELs <100bp within the coding region, and intergenic mutations (both SNPs and INDELs). For intergenic mutations the closest gene in bp to the SNP/INDEL is displayed.

General trends across all the pairs can be seen in Figure 3, SNPs were observed across pairs in genes involved in amino acid metabolism, DNA replication, metal transport/metabolism, sugar transport/metabolism and genes associated with mobile genetic elements and phages. Overall, most SNPs were found in genes related to prophages which contributed to a range of functions, but mostly within genes encoding phage endopeptidase, phage lysozyme and genes involved in the assembly of the phage tail. Synonymous SNPs were found in the genes involved in the synthesis of the amino acid glutamate in FS2240 and FS1654. Mutations in and around genes of the *lac* operon were found in FS2240 and FS2130 which included non-synonymous SNPs, INDELs and intergenic mutations. Synonymous SNPs and INDELs were found in the virulence gene *vgrG* in both FS1654 and FS2240.

FS2155 is an outlier with 516 SNPs and therefore the SNPs are displayed separately in Supplementary Table 5 . Non-synonymous SNPs of note were found in genes involved in sugar metabolism/transport, including genes of the *lac* operon (e.g. beta-galactosidase) and those associated with galactose, glucose, lactose, sucrose and mannose. Genes encoding transcriptional regulators of the *lac* operon were also found to have non-synonymous SNPs. Ten non-synonymous SNPs and 3 intergenic mutations were found in the *fec* genes encoding for the iron dicitrate transport system, with 1 non-synonymous mutation found in the ferrichrome-iron receptor gene. One of the non-synonymous SNPs in *fecA* was an A16T mutation predicted to not be tolerated in the protein structure and therefore is likely to affect protein function. Together this shows a complex picture of SNPs in a varied set of genes and gene classes contributing to, or because of, translocation to the blood.

### Insertions, deletions and mobile genetic element movement between highly related paired isolates

We investigated any discrepancies across the entire genome between the highly related pairs such as large insertions and deletions. The hybrid assemblies allowed us to compare contigs directly across pairs (see Supplementary Table 6) and a BLAST alignment of the blood and faecal isolates for each pair considered large insertions and deletions between and within contigs (Figure 4). All the highly related pairs, except FS1654 and FS2140, have 1 chromosome and between 0 and 8 plasmids, however it is worth noting that the term “plasmid” here is used to refer to any extra-chromosomal contig, and not all extra-chromosomal contigs are likely to be plasmids, as some are <5kb and contain no identifiable *rep* gene. Generally, across the pairs there is a loss or gain of phage genes and mobile genetic elements when translocating from the GI tract to the blood (see Supplementary Figure 2). The blood and faecal genomes of FS2155 have the largest disparities of all paired samples in Figure 4, with a 400kb disparity in chromosome size however this could be explained by the insertion of FC p1 into the chromosome (see Supplementary Figure 2). The connections in Figure 4 show that all 8 highly related pairs (from patients FS1386, FS1448, FS1654, FS2071, FS2112, FS2130, FS2240 and FS2258) have chromosomes which share greater than 99.99% sequence identity between blood and faecal isolates, providing further evidence that all these bloodstream isolates originated in the GI tract of their respective host.

**Figure 4:**
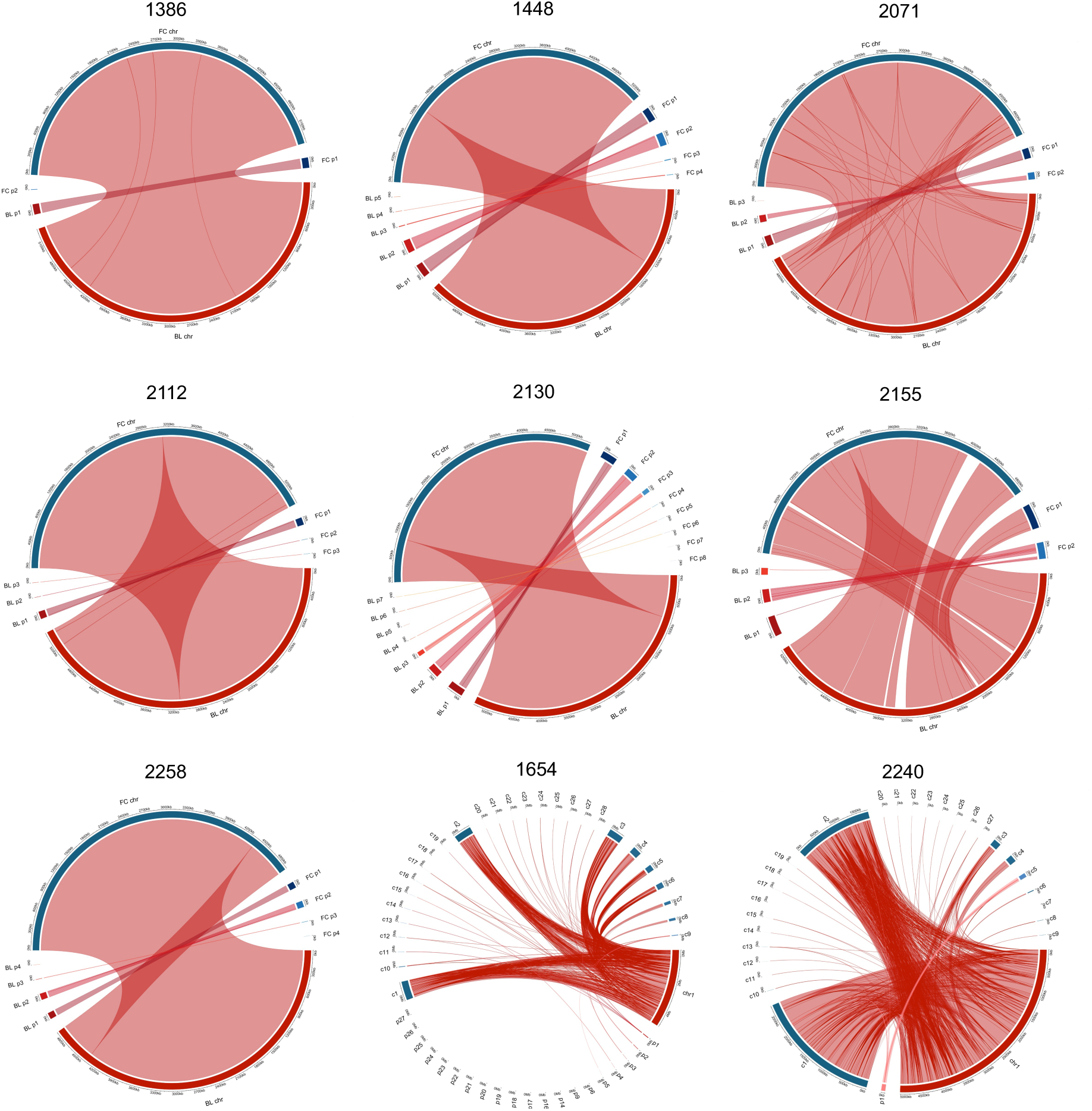
Whole genome comparison between the blood and faecal isolates for each highly related paired isolate. Connections shown are homologous regions which share greater than 99.9% sequence identity and are larger than 2kb for all pairs except FS1654 and FS2240 where no size limit was imposed (i.e. >0kb) due to the method of assembly. Each pair was aligned with BLASTn, blood isolate genome assemblies (red) were used as query sequence and faecal isolate genome assemblies (blue) were used as reference. Each genome is split into individual contigs, each with one chromosome and any extra-chromosomal contigs designated plasmids (even if a *rep* gene was absent). The exception is FS1654 and FS2240 where all contigs are numbered and not distinguished as chromosome or plasmid. FC, faecal isolates; BL, blood isolate; chr, chromosome; p, plasmid; c, contig.

### Hypermutator assay

We investigated whether FS2155 was a hypermutator strain which could explain why there is a difference in the MLST, a lower ANI and large genomic changes between blood and faecal isolates. There was no clear difference between FS2155 and the other isolates in their ability to become resistant to rifampicin, indicating that the differences in the FS2155 isolates are not due to it being a hypermutator strain under the laboratory conditions under which it was tested. (Supplementary Table 7).

### Virulence, pathogenicity and invasiveness determinant genes across all isolates

We screened all 26 isolates against the virulencefinder database to determine if there were certain genes which determined invasiveness/virulence in the translocation to the blood stream (Figure 5A). The *E. coli* and *K. pneumoniae* isolates can be easily distinguished by the absence or presence of certain genes. With the *E. coli*, all 8 isolates have similar genes which is expected as they are all part of phylogroup B2, the only difference is that FS2071 has 7 additional adhesion genes and lacks 2 genes, one involved in host defence evasion (*kpsT*) and another expressing a toxin (*vat*) which are found in the rest of the *E. coli*. With the *Klebsiella* spp., 61% of isolates (11/18) have the same set of virulence genes, these are all involved in iron-uptake including a set of *ybt* genes, *irp* and *fyuA*. The *ybt* genes, involved in the production of the yersiniabactin siderophore, are present in the paired isolates which likely translocated from the GI tract to the blood but not present in the unpaired isolates. This is also the case for the *irp2*, encoding an iron regulatory protein, and *fyuA*, encoding the yersiniabactin receptor. This could indicate that these *ybt*, *fyuA* and *irp2* genes are important for the translocation of *K. pneumoniae* into the blood. These genes are always found on the chromosome of these isolates and are co-located within the same biosynthetic gene cluster (Figure 5C). Of the rest of the *Klebsiella* isolates 22% (4/18) have only one virulence gene *yagZ*/*ecpA*, involved in adhesion, and 17% (3/18) had no detectable virulence gene from the virulence factor database.

**Figure 5:**
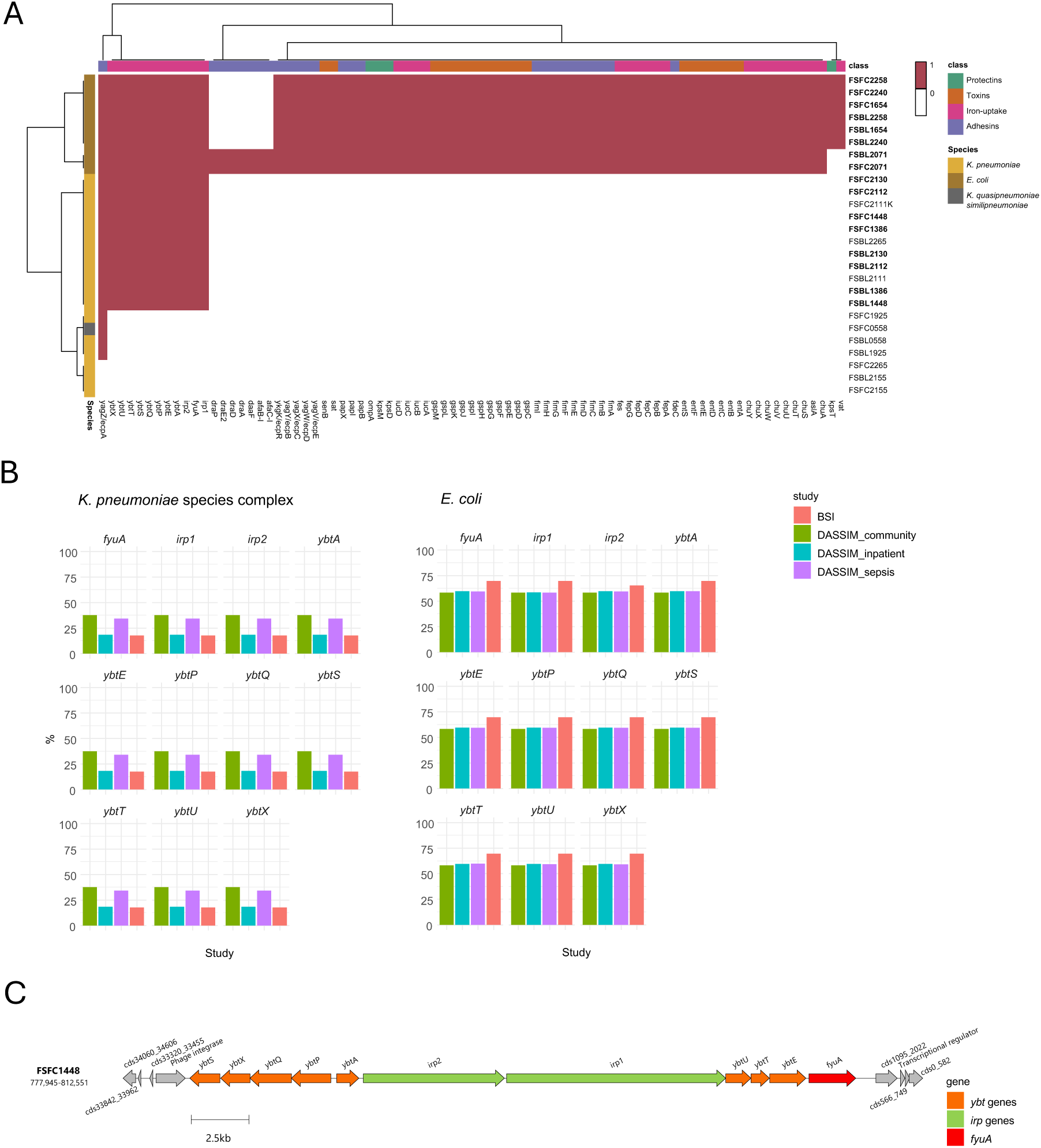
Virulence determinant genes in blood and faecal isolates across several datasets. A) Paired and unpaired isolates from this study (*n* = 26) were screened against the virulence factor database and the virulence genes present were visualised as a heatmap. Highly related pairs are in bold. The virulence genes were split into four classes. B) The percentage of the *ybt*, *fyuA* and *ipA* genes in two datasets, BSI and DASSIM, from Blantyre, Malawi containing 472 *E. coli* and 244 *K. pneumoniae* species complex isolates (KpSc). Out of the 472 *E. coli* 23 were BSI, 31 were DASSIM community, 334 were DASSIM sepsis and 84 were DASSIM inpatient. Out of the 244 KpSc 57 were BSI, 16 were DASSIM community, 138 were DASSIM sepsis and 33 were DASSIM inpatient. C) The genomic structure of the *ybt* biosynthetic gene cluster in the FSFC1448 genome.

To further investigate the association of the *ybt*, *fyuA* and *irp2* genes with BSI pathogenicity we analysed two datasets, BSI and DASSIM, containing 716 *E. coli* and *K. pneumoniae* species complex isolates (KpSc) from Blantyre, Malawi. The BSI dataset contained blood isolates from sepsis patients and the DASSIM dataset contained faecal samples from community patients, inpatients and sepsis patients. We determined the percentage of these isolates containing the *ybt, fyuA* and *ipr2* genes (Figure 5B) to see if they were overrepresented in blood stream isolates compared to faecal isolates. Sixty-nine percent of *E. coli* isolates from BSI dataset had all *ybt, fyuA* and *ipr2* genes, whereas in the DASSIM dataset, 58% were found in community patients, 59% found in inpatients and 59% found in sepsis patients. In this case the *ybt, fyuA* and *ipr2* genes were overrepresented in the bloodstream infection isolates. With the *K. pneumoniae* the picture is different, with the genes present in 18% of BSI isolates, and from the DASSIM faecal isolates the genes are present in 18% of inpatients, 34% of sepsis patients and 38% of community patients.

## Discussion

In this study we used comparative genomics with blood isolates and faecal samples taken from the same neonatal patients ^7,22^ to determine if the bacterial isolates translocated from the gut to the blood and what genomic changes, if any, accompanied this translocation event. Our methodology and study setting are novel in that we used in-depth comparisons of hybrid assembled genomes from the same patients in sub-Saharan Africa. Using MLST, ANI, and core genome phylogenetic trees we found that eight out of the thirteen pairs (4 *E. coli* and 4 *K. pneumoniae*) shared the same MLST, had an ANI above 99.99% and had a very small difference in branch length on the phylogenetic tree. This high relatedness between the pairs indicates that the 4 *E. coli* and 4 *K. pneumoniae* isolates likely translocated from the GI tract to the blood prior to the onset of blood stream infection and fever, a phenomena which has been previously reported from NICUs in the USA^9,11^. The 4 *E. coli* highly-related pairs (from patients FS1654, FS2071, FS2240 and FS2258) were all of the phylogroup B2 (Table 1) which has been associated with pathogenic clones, particularly extraintestinal pathogenic *E. coli* (ExPEC) ^10^ and has been indicative of urinary tract ^21,26^ and blood stream infection ^27^. The 4 *K. pneumoniae* highly related pairs each shared the same within-pair K-type, O-type and *wzi* gene type, yet these differed between different pairs.

The blood and faecal *K. pneumoniae* isolates from patient FS2155 shared the same cgMLST, K-type, O-type and *wzi* gene type but had a different MLST and a lower ANI of 99.78 between blood and faecal isolates. Further comparison of whole genome sequences (Figure 4) also indicated it is unlikely to be the same strain as large parts of the chromosome did not share >99.9% sequence identity which would be expected if the strain was the same and in fact with the eight highly related pairs, the entire chromosome shares 99.9% homology.

Seemingly, the young age of the patient (1 day old) and the short time of fever (see Supplementary Table 2) would not accommodate enough generations of *K. pneumoniae* to allow for large number for SNPs between the FS2155 blood and faecal isolates, however no isolates were found to have an elevated mutation rate based on the results of the rifampicin mutation assay (Supplementary Table 7). Alternatively, we may have just isolated different but related isolates from the faeces and the blood, or the selection procedure may have missed the blood isolate in the faecal sample when in fact it was present.

The ANI comparisons and phylogenetic tree showed there was high relatedness between isolates from different patients in the same hospital such as FS2111 and FS2112 from Amana and FS2258, FS1654 and FS2240 from Temeke (see Figure 2) indicating potential nosocomial transmission between patients. Nosocomial transmission between patients within NICUs in the USA has been indicated prior to the onset of sepsis in another study ^9^. It is difficult to apply ANI thresholds to isolate relatedness, however a recent study showed >95% indicate the same species, >99.5% indicate the same sequence type and >99.99% indicates the same strain ^28^. Our methodology also included SNP distances (see Figure 1) which provides a clearer distinction between closely related isolates. In genomes containing millions of base pairs, percentages can provide misleading results, for example the *E. coli* K-12 strain has a reference genome with a size of 4,639,221bp, 400 SNPs would still give an ANI of 99.99% but they may not be clonal. Therefore, going forward we recommend the use of both ANI and SNP distances to determine isolate relatedness within a cohort of compared strains.

The antibiotic resistance profiles did not differ between the highly related pairs. Previously it has been noted that the presence of ESBL– producing Gram-negative isolates in the faeces of hospitalized adults is associated with the subsequent onset of sepsis ^9^ and all our isolates that caused blood stream infections contained the common ESBL gene *bla*_CTX-M-15_ which confers resistance to 3^rd^ generation cephalosporins and is the most common ESBL gene found in *E. coli* isolates worldwide ^29^. This is likely due to the similarity in phenotypic antibiotic resistance profiles being a pre-requisite for selection of isolates in this study. Comparative genomics showed there were common SNPs across patient paired isolates in amino acid metabolism, DNA replication, metal transport/metabolism, sugar transport/metabolism and genes associated with mobile genetic elements and phages. INDELs were found in genes of the *lac* operon in FS2240 and FS2130 (Figure 3). INDEL mutations were also found in the *lac* operon and beta-galactosidase gene in FS2155 as well as genes involved in the metabolism of a wide set of sugars. *E. coli* can utilise a variety of sugars for growth in the gastrointestinal tract ^19^ and mutations in sugar metabolism/transport pathways have shown to have an effect on the colonisation of the enterohemorrhagic *E. coli* strain EDL933 in the intestine of mice ^30^. Since all the patients involved in this study were aged <19 days, the main sugar source for gut bacteria would be lactose from breast or formula milk, this could explain the mutations in the *lac* operon as it would be redundant following a change of carbohydrate substrate.

When comparing highly related pairs, a synonymous mutation was found in a copper resistance transcriptional regulator in FS1654, and in FS2155 ten non-synonymous mutations were found within the *fec* genes which encodes for the iron dicitrate transport system. This includes a A16T mutation in *fecA,* predicted to affect protein function. In a previous study a separate non-synonymous mutation was found in *fecD* when assessing *E. coli* moving from the gut to the urinary tract ^17^.

Most SNPs were found in genes related to phage/prophage (Figure 3), this supports the whole genome comparative analysis in Figure 4 and Supplementary Figure 2 which show that differences in the genomes were mainly accounted for by movement of mobile elements, including phage. It could be the case that the movement from the nutrient rich environment of the gut to the nutrient sparse environment of the blood stream causes a stress response in the cell which in turn leads to widespread movement of mobile genetic elements (MGEs) and the activation of prophages and subsequent mobilisation of phages ^31,32^. This activation of mobile genetic elements could be beneficial to bacteria, and the MGEs, allowing them to adapt to new environmental stresses by increasing genetic variability, as the movement of MGEs intracellularly can lead to gene knockouts or upregulation of mobilised genes when their genetic context changes ^33^. Paradoxically the activation of prophage to a phage would likely be deleterious for the bacterial host and therefore there could be a selection bottleneck for mutated phage genes in isolates successfully translocating between the gut and blood. In this case, the inactivation of genes involved in the replication of phages, and subsequent lysis, would be advantageous to the bacterium ^34–36^.

This widespread movement of MGEs, including phage, could also enable the acquisition of beneficial traits from other bacteria, however, while there is a high abundance of plasmid transfer between bacteria in the gut ^37^, there is unlikely to be high plasmid transfer in blood as there is a high amount of transience. During conjugation the cells must come into contact with one another and sustain the interaction long enough for genetic material to be exchanged ^38^ and previously it has been shown that conjugation occurs at a higher rate in fixed conditions such as biofilms rather than liquid culture ^39–41^.

We also looked to see if there were any genes in our blood isolates that were indicative of virulence, pathogenicity and invasiveness by screening all isolates against the virulence factor database (Figure 5A). All *E. coli* isolates had the same virulence genes except the ones from FS2071 which had 7 additional adhesion genes and lacked 2 genes when compared to the others. The *Klebsiella* spp. had more variation in virulence genes, with 61%, including all the highly related pairs, having 11 genes involved in iron-uptake, and the rest (39%) only having 1 or 0 virulence genes. The additional 11 genes were variants of *ybt*, *irp* and *fyu* genes which are all involved in iron uptake and metabolism. The *ybt* gene encodes for the yersiniabactin siderophore which, unlike the enterobactin siderophore, can escape the host innate immune system protein sideocalin, and is associated with pathogenicity in *E. coli* and *K. pneumoniae* ^42^. This could indicate that the yersiniabactin siderophore is key for the translocation of *K. pneumoniae* into the blood since all the paired isolates had the yersiniabactin genes and the unpaired isolates did not contain these genes. These unpaired isolates unintentionally provide a background of faecal isolates that were not confirmed to translocate to the blood stream and therefore show the potential difference in genes between intestinal *Klebsiella* and extra-intestinal pathogenic *Klebsiella*. We tested this further by including two datasets from Blantyre, Malawi (BSI and DASSIM) to compare the presence of the *ybt*, *iprA* and *fyuA* genes in 716 blood and faecal isolates and see if the genes were overrepresented in the bloodstream isolates. While there was no clear distinction between blood and faecal *Klebsiella* isolates, in *E. coli* isolates there was a higher abundance of these genes in the BSI isolates when compared to faecal carriage isolates from DASSIM. Together, this indicates that important genes required for invasion into and / or persistence in the blood were already present in the gut isolates. This opens the possibility for diagnostic interventions, for example, certain patient groups, such as very low birthweight neonates, could have their stool screened by PCR for the presence of the *ybt*, *fyuA* and *irp2* genes to determine if they are predisposed to bloodstream infections. This would require further investigation, however if successful this targeted intervention could contribute to the prevention of BSIs in neonates.

Although this study expands our understanding of the transition of Enterobacterales from the gut to the bloodstream, it is not without limitations. We selected our isolates using single colony picks, therefore we may have missed some highly related paired isolates at the selection stage. Not all the originally paired isolates could be found and sequenced, therefore only 26 out of the original 32 isolates could be sequenced as pairs. By not sequencing these we may have lost information about genes or SNPs involved in bacterial translocation to the blood and we cannot predict how this may have biased the results. Further to this, while most of the genomes were assembled with the same hybrid assembly method (hybracter), due to a lack of short read sequences two were assembled with flye and due to poor long read quality two were assembled with unicycler. While this did not affect sequence typing, ANI, SNP analysis or screening for the absence of presence of genes it did have an impact on the whole genome comparison between the blood and faecal isolates (Figure 4).

To conclude, we have shown that 8 isolates (4 *E. coli* and 4 *K. pneumoniae*) likely moved from the gut to the blood in neonatal patients with blood stream infections. The changes that marked this transition varied between the isolate pairs but were mainly found in genes associated with mobile genetic elements and phages. It is possible that most of the genes required for invasion into, or persistence in, the blood were already present in the gut isolates as a set of important iron uptake genes were shared across blood and faecal isolates within each species. This analysis also highlighted the potential importance of the yersiniabactin siderophore in BSI caused by both *Klebsiella* spp. and *E. coli*. These findings expand our understanding of BSI pathogenesis and provide potential genetic markers which could be utilised in targeted interventions to prevent future BSI infections in neonates.

## Methods

### Sample collection and bacterial isolation

Enrolment of participants, collection of blood samples and bacterial isolation was carried out as described previously ^7^. The faecal samples were collected as rectal swabs from each patient and bacteria was isolated as previously described ^22^. The strains were sent to the Liverpool School of Tropical Medicine, UK and stored at −80°C until the DNA was extracted and sequenced.

### Whole genome sequencing

Short read sequencing was carried out by MicrobesNG (Birmingham, UK). Briefly, the DNA was extracted from the blood and faecal bacterial strains and prepared into DNA libraries using the Nextera XT Library Prep Kit (Illumina, California, USA). The short-read sequencing was carried out using an Illumina machine (Hiseq X10 platform) according to the 250bp paired end protocol.

For long read sequencing, DNA was extracted from blood isolates with the Fire Monkey Weight High molecular weight (HMW) genomic (gDNA) DNA Extraction Kit (Revolugen, UK) and the faecal isolates with the Wizard HMW DNA extraction kit (Promega, Wisconsin, USA). Library prep was carried out using the SQK-NBD114.24 Ligation Sequencing and Native Barcoding Kit according to the manufacturers protocol (Oxford Nanopore Technologies, Oxford, UK). Sequencing was carried out using a FLO-MIN114 (R10.4.1) flow cell (Oxford Nanopore Technologies, Oxford, UK) on a MinION Mk1B sequencer, running for 72 hr at a translocation speed of 400bp/s. Data acquisition used the MinKNOW software (v22.08.9).

### Hybrid genome assembly

The raw data (fast5) was basecalled using guppy v6.4.2 with the super accuracy (sup) model specific to the flow cell (R10.4.1), motor protein (E8.2) and translocation speed (400bp/s). Read quality was assessed with Nanoplot v1.38.1 ^43^. The basecalled files were demultiplexed the guppy_barcoder from Guppy v6.3.8 with the SQK-446NBD114-24 kit. The reads were assembled into *de novo* genome assemblies using either hybracter v0.6.0 ^44^, unicycler v0.4.8 ^45^ or flye v2.9.2 ^46^ (see Supplementary Table 1). The majority were assembled with hybracter (*n* = 22), FSFC1448 and FSBL1448 were assembled with flye as no short read match could be found and FSFC1654 and FSFC2240 were assembled with unicycler due to the short reads being better quality than the long reads. Hybracter is an automated pipeline for long-read first hybrid assembly. This workflow runs initial quality control, assembles the genome from long reads with flye ^46^ and plasmids with plassembler ^47^, polishes with long reads using medaka (https://github.com/nanoporetech/medaka), polishes with short reads using polypolish ^48^ and pypolca (https://github.com/gbouras13/pypolca) and returns an output of assembly files and statistics. This workflow has been shown to be more accurate than the existing gold-standard hybrid assembly software ^44^. Assemblies were assessed for quality with seqkit ^49^ (see Supplementary Table 1) and visualised with Bandage v0.8.1 ^50^.

### Genomic data analysis and phylogenetics

The assembled genomes were annotated using RAST ^51^ and screened against the Resfinder ^52^, virulence factor database (VFDB) ^53^ and MEGARes ^54^ databases using abricate v1.0.1 (https://github.com/tseemann/abricate). Average Nucleotide Identities (ANI) between genomes were determined using JSpeciesWS v4.1.1 ^55^ and SNP distances were determined using snippy v4.6.3 and snp-dists v0.8.2 with the *K. pneumoniae* FSFC1386 isolate as reference. The heatmap showing both ANI and SNP distances between isolates was visualised with the pandas v 2.2.3, matplotlib v 3.9.4, seaborn v 0.13.2 and numpy v 2.2.0 packages in python v3.10.12. Sequence typing used multilocus sequence types (MLST) and core genome MLST (cgMLST) schemes as part of the Pathogenwatch platform ^56^. The MLST database from Pasteur Institute, France was used for *Klebsiella* species (https://bigsdb.pasteur.fr/klebsiella/) and from Warwick University, UK for *E. coli* (http://mlst.warwick.ac.uk/mlst/dbs/Ecoli). The cgMLST database from Pasteur Institute, France was used for *Klebsiella* species (https://bigsdb.pasteur.fr/klebsiella/cgmlst-lincodes/). O- and K-types for *Klebsiella* species were predicted with Kaptive ^24^ along with *wzi* capsular gene type ^57^. For *E. coli* phylogroups were assigned using ClermonTyping ^25^ which is an *in silico* typing method based on the quadruplex PCR of the *arpA*, *chuA*, *yjaA* and *TspE4.C2* genes ^58^.

A core genome phylogenetic tree of 17 *K. pneumoniae* and 8 *E. coli* isolates was built (*n* = 25), the *K. quasipneumoniae* isolate FSFC0558 was omitted from this analysis so that intra-species relationships could be analysed. The isolates were aligned to species-specific reference genomes using snippy v 4.3.6, with the *Klebsiella pneumoniae* subsp. *pneumoniae* HS11286 genome (GCF_000240185.1) used as reference for *Klebsiella pneumoniae* isolates and the *Escherichia coli* str. K-12 substr. MG1655 genome (GCF_000005845.2) used as reference for *E. coli* isolates. The core SNP alignments were then combined using snippy-core. To retain only vertically inherited SNPs gubbins v2.3.4 was used to remove any recombinant regions from the alignment. A SNP-only alignment was extracted using snp-sites v2.5.1 with the -c option to select for A,C,G and T only. This was passed to IQ-tree v 1.6.1 which was used to construct a maximum-likelihood phylogenetic tree. The model was determined using the ModelFinder function of IQ-tree (-m MF) which calculated TVM+F+ASC+R2 as the best model for *K. pneumoniae* alignments and TVMe+ASC for *E. coli* alignments. The model was run using ultrafast 1,000 bootstrap replicates (-bb 1000) and keeping identical data (-keep-ident). The phylogenetic tree and associated metadata was visualised using Interactive Tree Of Life (iTOL) v6 ^59^. The timelines were plotted from the admission metadata using dplyr v1.1.4 and ggplot v3.5.1.

Breseq v0.38.3 ^60^ was used to compare the RAST annotated genomes from the faecal isolates against the long reads from the blood isolates, with the predicted mutations and unassigned missing coverage (MC) analysed from the output. The genes of interest were classified into clusters of orthologous genes (COGs) ^61^ using eggNOG v5.0 ^62^, separated according to their mutations, and visualised with the pheatmap v1.0.12 package in R v4.3.1. Genes which were categorised as “Function unknown” within the COG framework were further investigated for putative functions leading to an additional classification termed “class” inferred from gene annotations and BLAST ^63^ searches against similar genes. We used SIFT ^64^ to determine the predicted effect of genetic mutations on their protein function.

To determine the whole genome comparison between pairs the hybrid assembly from blood and faecal isolates were compared with BLASTn ^63^ with the faecal isolate as query compared against a database made from the blood isolate genome. The BLAST output was wrangled with dplyr v1.1.4 to visualise as a circos diagram with the “genomic initialize” function of the circlize v0.4.16 package in R v4.3.1. Connections plotted were homologous regions which shared greater than 99.9% sequence identity and were larger than 2kb except for FS1654 and FS1654 where connections were homologous regions with > 99.9% sequence identity with no threshold limit on basepair size.

The 26 isolates from this study were screened against the virulence factor database (VFDB) ^53^ using abricate v1.0.1 (https://github.com/tseemann/abricate), the output was plotted as a heatmap with the pheatmap v1.0.12 package in R v4.3.1. Two datasets, BSI and DASSIM, were analysed from Blantyre, Malawi to determine percentage of iron uptake virulence genes (*ybt*, *fyuA* and *ipA*) in blood and faecal isolates from another country in sub-Saharan Africa. Raw sequence read data was downloaded from the European Nucleotide Archive from the following projects: PRJEB8265, PRJEB28522, PRJEB26677, and PRJEB36486 ^65–67^.

The BSI dataset contains isolates from blood, cerebral spinal fluid (CSF) and rectal swabs, collected as part of routine BSI surveillance, only the blood isolates were used for this analysis^65,66^. The DASSIM dataset involved stool sampling and ESBL-E isolate selection from patients with sepsis^67^. The sequence reads were quality controlled, trimmed and assembled as previously described ^68^. 716 isolates (472 *E. coli* and 244 *Klebsiella* spp.) were screened against the VFDB ^53^ using abricate v1.0.1. The data were separated by dataset and group (DASSIM_sepsis, DASSIM_inpatient, DASSIM_community, and BSI) and the percentage of occurrences were calculated by the number of gene occurrences divided by the total number in each group per species. Calculations were performed using the “summarize” function in dplyr v1.1.4 and the barplots were visualised with ggplot v3.5.1. The genomic structure of the *ybt* biosynthetic gene cluster from the RAST^51^ annotated FSFC1448 genome was visualised using clinker v0.0.31 ^69^.

### Calculation of mutation frequency

A rifampicin resistance assay was used to determine the mutation frequency for seven faecal isolates. The colonies were incubated at 37°C in LB broth for 18hrs, the cultures were normalised to an OD_600_ of 0.1, then incubated at 37°C until the OD_600_ reached 0.6-0.8. The cultures were centrifuged at 4,000 rpm to pellet the cells then resuspended in 1ml of PBS from this resuspended solution, 100ml was diluted in PBS to 10^-6^ and plated on LB agar plates and 100ml was plated on LB agar containing rifampicin (20µg/ml). Plates were incubated at 37°C for 16hrs. CFU/ml was determined from colony counts of the total cell count plates. Mutation frequency was calculated by dividing the number of rifampicin-resistant colonies by the total number of viable cells plated (CFU/mL).

## Data availability

Reads from all isolates sequenced as part of this study have been submitted to the Sequence Read Archive (SRA) of the National Centre for Biotechnology Information (NCBI) under the project ID PRJNA1254181. Supplementary Figures 1-2 and Supplementary Tables 1-7 are accessible in the supplementary material of this manuscript.

## Code availability

All code to reproduce this analysis is publicly available through GitHub: https://rngoodman.github.io/blood-faecal-genomic-comparison.

## Acknowledgements

R.N.G and S.J.M. are supported by funding from the Medical Research Council (MRC), Biotechnology and Biological Sciences Research Council (BBSRC) and Natural Environmental Research Council (NERC), which are all Councils of UK Research and Innovation (Grant no. MR/W030578/1) and the Research Council of Norway (grant NFR333432) under the umbrella of the JPIAMR - Joint Programming Initiative on Antimicrobial Resistance via the STRESST project.

## Contributions

Conceptualization: R.N.G., S.J.M., B.B., N.L., A.P.R.; formal analysis: R.N.G., S.J.M., and A.P.R.; funding acquisition: A.P.R., N.L.; investigation: R.N.G., S.J.M., I.M., A.K., J.M., U.O.K., S.A., B.B., N.L. and A.P.R.; resources: S.J.M., J.M., U.O.K., S.A., B.B., N.L.; software: R.N.G.; methodology: R.N.G.; project administration: R.N.G., S.J.M., B.B., N.L., A.P.R.; supervision: S.J.M., B.B., N.L., A.P.R.; writing – original draft: R.N.G.; writing – review and editing: R.N.G., S.J.M, A.K., S.A., B.B., N.L., A.P.R.

## Ethics declarations

The original sample collection was part of a study approved by the Senate Research and Publications Committee of Muhimbili University of Health and Allied Sciences, National Institute of Medical Research, Tanzania and the Regional Committee for Medical and Health Research Ethics (REK), Norway.

