## Supplementary Tables 1, 2, 4, 6 and 7; Supplementary Figures 1 and 2 for "Genomic comparison of highly related pairs of *E. coli* and *K. pneumoniae* isolated from faeces and blood of the same neonatal patients hospitalized with fever in Dar es Salaam, Tanzania"

### Supplementary data

**Supplementary Table 1: QC of assembled genomes used in analysis.** The reads were assembled with either hybracter (*n*=22), unicycler (*n*=2) or flye (*n*=2).

| Sample | Assembly method | file format | type | number of seqs | sum length | min length | average length | max length | Q1 | Q2 | Q3 | N50 | GC(%) |
|---|---|---|---|---|---|---|---|---|---|---|---|---|---|
| FSBL0558 | hybracter | FASTA | DNA | 3 | 5,544,499 | 4,259 | 1,848,166.30 | 5,390,483 | 77,008 | 149,757 | 2,770,120 | 5,390,483 | 57.13 |
| FSBL1386 | hybracter | FASTA | DNA | 2 | 5,484,582 | 143,420 | 2,742,291 | 5,341,162 | 143,420 | 2,742,291 | 5,341,162 | 5,341,162 | 57.33 |
| FSBL1448 | flye | FASTA | DNA | 6 | 5,785,608 | 5,763 | 964,268 | 5,328,540 | 5,913 | 112,375 | 220,642 | 5,328,540 | 56.71 |
| FSBL1654 | hybracter | FASTA | DNA | 28 | 5,065,408 | 106 | 180,907.40 | 4,961,302 | 125 | 291.5 | 1,533.50 | 4,961,302 | 50.65 |
| FSBL1925 | hybracter | FASTA | DNA | 6 | 5,479,388 | 3,676 | 913,231.30 | 5,242,001 | 4,907 | 19,816.50 | 189,171 | 5,242,001 | 57.31 |
| FSBL2071 | hybracter | FASTA | DNA | 4 | 5,296,687 | 1,378 | 1,324,171.80 | 5,052,976 | 49,500 | 121,166.50 | 2,598,843.50 | 5,052,976 | 50.72 |
| FSBL2111 | hybracter | FASTA | DNA | 4 | 5,609,861 | 3,574 | 1,402,465.30 | 5,488,634 | 3,794 | 58,826.50 | 2,801,136.50 | 5,488,634 | 57.19 |
| FSBL2112 | hybracter | FASTA | DNA | 4 | 5,609,862 | 3,574 | 1,402,465.50 | 5,488,634 | 3,794 | 58,827 | 2,801,137 | 5,488,634 | 57.19 |
| FSBL2130 | hybracter | FASTA | DNA | 8 | 5,895,662 | 3,301 | 736,957.80 | 5,278,286 | 3,392.50 | 42,972.50 | 260,672.50 | 5,278,286 | 56.7 |
| FSBL2155 | hybracter | FASTA | DNA | 4 | 5,914,917 | 100,163 | 1,478,729.30 | 5,324,353 | 146,693 | 245,200.50 | 2,810,765.50 | 5,324,353 | 56.68 |
| FSBL2240 | hybracter | FASTA | DNA | 4 | 5,108,105 | 2,125 | 1,277,026.30 | 5,009,705 | 3,099.50 | 48,137.50 | 2,550,953 | 5,009,705 | 50.65 |
| FSBL2258 | hybracter | FASTA | DNA | 5 | 5,129,516 | 2,125 | 1,025,903.20 | 4,939,041 | 5,164 | 90,985 | 92,201 | 4,939,041 | 50.62 |
| FSBL2265 | hybracter | FASTA | DNA | 9 | 5,997,171 | 3,301 | 666,352.30 | 5,278,317 | 3,468 | 75,683 | 226,098 | 5,278,317 | 56.66 |
| FSFC0558 | hybracter | FASTA | DNA | 2 | 5,269,471 | 166,709 | 2,634,735.50 | 5,102,762 | 166,709 | 2,634,735.50 | 5,102,762 | 5,102,762 | 58.02 |
| FSFC1386 | hybracter | FASTA | DNA | 3 | 5,490,501 | 6,415 | 1,830,167 | 5,340,663 | 74,919 | 143,423 | 2,742,043 | 5,340,663 | 57.31 |
| FSFC1448 | flye | FASTA | DNA | 5 | 5,786,169 | 10,122 | 1,157,233.80 | 5,328,480 | 14,596 | 212,329 | 220,642 | 5,328,480 | 56.71 |
| FSFC1654 | unicycler | FASTA | DNA | 28 | 5,042,889 | 131 | 180,103.20 | 1,190,061 | 843.5 | 9,908.50 | 172,451 | 933,194 | 50.64 |
| FSFC1925 | hybracter | FASTA | DNA | 4 | 5,684,240 | 63,275 | 1,421,060 | 5,290,046 | 87,407 | 165,459.50 | 2,754,713 | 5,290,046 | 57.03 |
| FSFC2071 | hybracter | FASTA | DNA | 3 | 5,295,994 | 98,286 | 1,765,331.30 | 5,052,997 | 121,498.50 | 144,711 | 2,598,854 | 5,052,997 | 50.72 |
| FSFC2111K | hybracter | FASTA | DNA | 2 | 5,593,489 | 229,590 | 2,796,744.50 | 5,363,899 | 229,590 | 2,796,744.50 | 5,363,899 | 5,363,899 | 57.07 |
| FSFC2112 | hybracter | FASTA | DNA | 4 | 5,608,700 | 3,574 | 1,402,175 | 5,488,573 | 3,794 | 58,276.50 | 2,800,556 | 5,488,573 | 57.19 |
| FSFC2130 | hybracter | FASTA | DNA | 9 | 5,885,363 | 873 | 653,929.20 | 5,278,286 | 3,301 | 4,761 | 224,982 | 5,278,286 | 56.71 |
| FSFC2155 | hybracter | FASTA | DNA | 3 | 5,537,134 | 244,643 | 1,845,711.30 | 4,924,353 | 306,390.50 | 368,138 | 2,646,245.50 | 4,924,353 | 57.17 |
| FSFC2240 | unicycler | FASTA | DNA | 27 | 5,084,135 | 131 | 188,301.30 | 2,839,153 | 328 | 1,225 | 11,260 | 2,839,153 | 50.64 |
| FSFC2258 | hybracter | FASTA | DNA | 5 | 5,129,448 | 2,125 | 1,025,889.60 | 4,939,014 | 5,164 | 90,944 | 92,201 | 4,939,014 | 50.62 |
| FSFC2265 | hybracter | FASTA | DNA | 27 | 5,645,840 | 232 | 209,105.20 | 5,172,443 | 1,970.50 | 4,761 | 22,757.50 | 5,172,443 | 57 |

**Supplementary Table 2: List of clinical metadata associated with each blood and faecal paired isolates.** The ID number refers to individual patients.

| Isolate | Species | Sample source: | ID number | Hospital | Year | Age on admission | Duration | Antibiotics before admission | Antibiotics before culture | Antibiotics |
|---|---|---|---|---|---|---|---|---|---|---|
| FSFC0558 | *K. quasipneumoniae similipneumoniae* | Faecal | FS0558 | MNH | 2017 | 1 day (<24hrs) | (<24hrs) | No | Yes | Amoxycillin+genta |
| FSFC1386 | *K. pneumoniae* | Faecal | FS1386 | Temeke | 2018 | 2 days | 2 | No | Yes | Amoxycillin+genta |
| FSFC1448 | *K. pneumoniae* | Faecal | FS1448 | Temeke | 2018 | 2 days | 2 | No | Yes | Amoxycillin+genta |
| FSFC1654 | *E. coli* | Faecal | FS1654 | Temeke | 2018 | 1 day | 1 day | No | Yes | Amoxycillin+genta |
| FSFC1925 | *K. pneumoniae* | Faecal | FS1925 | Amana | 2018 | 3 days | 2 days | No | Yes | Amoxycillin /then Ceftriaxone +genta |
| FSFC2071 | *E. coli* | Faecal | FS2071 | Temeke | 2018 | 4 days | 2 days | No | Yes | Amoxycillin+genta |
| FSFC2111K | *K. pneumoniae* | Faecal | FS2111 | Amana | 2018 | 2 days | 1 day | No | Yes | Amoxycillin /then Ceftriaxone +genta |
| FSFC2112 | *K. pneumoniae* | Faecal | FS2112 | Amana | 2018 | 3 days | 2 days | No | Yes | Amoxycillin /then Ceftriaxone +genta |
| FSFC2130 | *K. pneumoniae* | Faecal | FS2130 | Amana | 2018 | 3 days | 2 days | No | Yes | Amoxycillin /then Ceftriaxone +genta |
| FSFC2155 | *K. pneumoniae* | Faecal | FS2155 | Amana | 2018 | 1 day | 1 day | No | Yes | Amoxycillin+genta |
| FSFC2240 | *E. coli* | Faecal | FS2240 | Temeke | 2018 | 16 days | 2 days | No | Yes | Amoxycillin+genta |
| FSFC2258 | *E. coli* | Faecal | FS2258 | Temeke | 2018 | 1 day | 1 day | No | Yes | Amoxycillin+genta |
| FSFC2265 | *K. pneumoniae* | Faecal | FS2265 | Temeke | 2018 | 19 days | 3 days | No | Yes | Amoxycillin+genta |
| FSBL0558 | *K. pneumoniae* | Blood | FS0558 | MNH | 2018 | 1 day (<24hrs) | (<24hrs) | No | Yes | Amoxycillin+genta |
| FSBL1386 | *K. pneumoniae* | Blood | FS1386 | Temeke | 2018 | 2 days | 2 | No | Yes | Amoxycillin+genta |
| FSBL1448 | *K. pneumoniae* | Blood | FS1448 | Temeke | 2018 | 2 days | 2 | No | Yes | Amoxycillin+genta |
| FSBL1654 | *E. coli* | Blood | FS1654 | Temeke | 2018 | 1 day | 1 day | No | Yes | Amoxycillin+genta |
| FSBL1925 | *K. pneumoniae* | Blood | FS1925 | Amana | 2018 | 3 days | 2 days | No | Yes | Amoxycillin /then Ceftriaxone +genta |
| FSBL2071 | *E. coli* | Blood | FS2071 | Temeke | 2018 | 4 days | 2 days | No | Yes | Amoxycillin+genta |
| FSBL2111 | *K. pneumoniae* | Blood | FS2111 | Amana | 2018 | 2 days | 1 day | No | Yes | Amoxycillin /then Ceftriaxone +genta |
| FSBL2112 | *K. pneumoniae* | Blood | FS2112 | Amana | 2018 | 3 days | 2 days | No | Yes | Amoxycillin /then Ceftriaxone +genta |
| FSBL2130 | *K. pneumoniae* | Blood | FS2130 | Amana | 2018 | 3 days | 2 days | No | Yes | Amoxycillin /then Ceftriaxone +genta |
| FSBL2155 | *K. pneumoniae* | Blood | FS2155 | Amana | 2018 | 1 day | 1 day | No | Yes | Amoxycillin+genta |
| FSBL2240 | *E. coli* | Blood | FS2240 | Temeke | 2018 | 16 days | 2 days | No | Yes | Amoxycillin+genta |
| FSBL2258 | *E. coli* | Blood | FS2258 | Temeke | 2018 | 1 day | 1 day | No | Yes | Amoxycillin+genta |
| FSBL2265 | *K. pneumoniae* | Blood | FS2265 | Temeke | 2018 | 19 days | 3 days | No | Yes | Amoxycillin+genta |

**Supplementary Table 3: Resfinder, Virulence factor and MEGARES database screening of 8 highly related pairs.** All pairs were hybrid assembled with hybracter except FS1448 where a flye assembly of long reads is used and FS1654 and FS2240 where a unicycler hybrid assembly was used.

See separate Excel spreadsheet:

*Supplementary Table 3 - Resfinder, Virulence factor and MEGARES database screening of 8 highly related pairs.xlsx*

**Supplementary Table 4: SNP and small INDEL (<100bp) frequencies from breseq whole genomic comparison.**

| Breseq SNP and INDELs | |
|---|---|
| Sample | Frequency |
| FS1386 | 1 |
| FS1448 | 8 |
| FS1654 | 10 |
| FS2071 | 97 |
| FS2112 | 4 |
| FS2130 | 6 |
| FS2258 | 23 |
| FS2240 | 34 |
| FS2155 | 516 |

| | |
|---|---|
| Average | 77.66 |
| Min | 1 |
| Max | 516 |
| Median | 10 |
| Standard deviation | 167.06 |

**Supplementary Table 5: Breseq comparisons of FSFC2155 hybrid assembly against FSBL2155 long reads.** 516 SNPs in 289 genes are displayed grouped by gene classification.

See separate Excel spreadsheet:

*Supplementary Table 5 - Breseq comparisons of FSFC2155 hybrid assembly against FSBL2155 long reads.xlsx*

**Supplementary Table 6: Genome assembly statistics by contigs across highly related pairs with hybrid assemblies.** Isolates from FS1654 and FS2240, although highly related pairs were omitted as their assemblies have > 25 contigs. Isolates from FS2155 are included for reference although not classed as a highly related pair. Difference is calculated by subtracting contig sizes of blood isolate by contig sizes of faecal isolate.

| Isolate name | Sample source | Isolate name | Contig name | length (bp) | GC (%) | Circular | Sample source | Isolate name | Contig name | length | GC | Circular | Difference in blood relative to fecal (+/- |
|---|---|---|---|---|---|---|---|---|---|---|---|---|---|
| FS1386 | Fecal | FSFC1386 | chromosome | 5340663 | 57.48 | FALSE | Blood | FSBL1386 | chromosome | 5341162 | 57.48 | TRUE | 499 |
| | | | plasmid 1 | 143423 | 51.87 | TRUE | | | plasmid 1 | 143420 | 51.87 | TRUE | -3 |
| | | | Plasmid 2 | 6415 | 40.12 | TRUE | | | | | | | -6415 |
| FS1448* | | FSFC1448 | chromosome | 5328480 | 57.4 | FALSE | | FSBL1448 | chromosome | 5328540 | 57.4 | FALSE | 60 |
| | | | plasmid 1 | 220642 | 52.08 | FALSE | | | plasmid 1 | 220642 | 52.08 | FALSE | 0 |
| | | | plasmid 2 | 212329 | 45.19 | FALSE | | | plasmid 2 | 212332 | 45.19 | FALSE | 3 |
| | | | plasmid 3 | 14596 | 43.45 | FALSE | | | plasmid 3 | 12418 | 52.48 | FALSE | -2178 |
| | | | plasmid 4 | 10122 | 53.39 | FALSE | | | plasmid 4 | 5913 | 43.62 | FALSE | -4209 |
| | | | | | | | | | plasmid 5 | 5763 | 42.11 | FALSE | 5763 |
| FS2071 | | FSFC2071 | Chromsome | 5052997 | 50.72 | TRUE | | FSBL2071 | chromosome | 5052976 | 50.72 | TRUE | -21 |
| | | | plasmid 1 | 144711 | 52.6 | TRUE | | | plasmid 1 | 144711 | 52.6 | TRUE | 0 |
| | | | Plasmid 2 | 98286 | 47.91 | TRUE | | | plasmid 2 | 97622 | 47.92 | TRUE | -664 |
| | | | | | | | | | plasmid 3 | 1378 | 47.39 | FALSE | 1378 |
| FS2112 | | FSFC2112 | chromosome | 5488573 | 57.3 | TRUE | | FSBL2112 | chromosome | 5488634 | 57.3 | TRUE | 61 |
| | | | plasmid 1 | 112539 | 52.78 | TRUE | | | plasmid 1 | 113640 | 52.77 | TRUE | 1101 |
| | | | plasmid 2 | 4014 | 43.72 | TRUE | | | plasmid 2 | 4014 | 43.72 | TRUE | 0 |
| | | | plasmid 3 | 3574 | 43.84 | TRUE | | | plasmid 3 | 3574 | 43.84 | TRUE | 0 |
| FS2130 | | FSFC2130 | chromosome | 5278286 | 57.51 | TRUE | | FSBL2130 | chromosome | 5278286 | 57.51 | TRUE | 0 |
| | | | plasmid 1 | 287690 | 47.06 | FALSE | | | plasmid 1 | 296347 | 47.17 | TRUE | 8657 |
| | | | plasmid 2 | 224982 | 51.98 | TRUE | | | plasmid 2 | 224998 | 51.98 | TRUE | 16 |
| | | | plasmid 3 | 81179 | 54 | TRUE | | | plasmid 3 | 81184 | 54.05 | TRUE | 5 |
| | | | plasmid 4 | 4761 | 44.42 | TRUE | | | plasmid 4 | 4761 | 44.42 | TRUE | 0 |
| | | | plasmid 5 | 3317 | 46.28 | TRUE | | | plasmid 5 | 3468 | 45.13 | TRUE | 151 |
| | | | plasmid 6 | 3301 | 43.02 | TRUE | | | plasmid 6 | 3317 | 46.28 | TRUE | 16 |
| | | | plasmid 7 | 974 | 40.76 | FALSE | | | plasmid 7 | 3301 | 43.02 | TRUE | 2327 |
| | | | plasmid 8 | 873 | 43.64 | FALSE | | | | | | | |
| FS2155 | | FSFC2155 | chromosome | 4924353 | 57.28 | FALSE | | FSBL2155 | chromosome | 5324353 | 57.42 | TRUE | 400000 |
| | | | plasmid 1 | 368138 | 58.77 | FALSE | | | plasmid 1 | 297178 | 47.19 | TRUE | -70960 |
| | | | plasmid 2 | 244643 | 52.58 | FALSE | | | plasmid 2 | 193223 | 52.01 | TRUE | -51420 |
| | | | | | | | | | plasmid 3 | 100163 | 54.24 | TRUE | 100163 |
| FS2258 | | FSFC2258 | chromosome | 4939014 | 50.62 | TRUE | | FSBL2258 | chromosome | 4939041 | 50.62 | TRUE | 27 |
| | | | plasmid 1 | 92201 | 51.41 | TRUE | | | plasmid 1 | 92201 | 51.41 | TRUE | 0 |
| | | | plasmid 2 | 90944 | 50.12 | TRUE | | | plasmid 2 | 90985 | 50.11 | TRUE | 41 |
| | | | plasmid 3 | 5164 | 47.52 | TRUE | | | plasmid 3 | 5164 | 47.52 | TRUE | 0 |
| | | | plasmid 4 | 2125 | 47.11 | TRUE | | | plasmid 4 | 2125 | 47.11 | TRUE | 0 |

**Supplementary Table 7: Rifampicin mutation assay.** Mutation frequency was calculated by dividing the number of rifampicin-resistant colonies by the total number of viable cells plated.

| Sample | CFU $ml^{-1}$ | CFU $100\mu l^{-1}$ | Growth on LB+RIF (20µg/ml) after 16hrs | | | | Mutation Frequency |
|---|---|---|---|---|---|---|---|
| | | | 1 | 2 | 3 | Average | |
| FS1386 | $1.74 \times 10^{7}$ | $1.74 \times 10^{6}$ | 0 | 0 | 0 | 0 | 0 |
| FS1448 | $1.69 \times 10^{7}$ | $1.69 \times 10^{6}$ | 0 | 1 | 1 | 0.66666667 | $3.94 \times 10^{-7}$ |
| FS2071 | $1.56 \times 10^{7}$ | $1.56 \times 10^{6}$ | 0 | 0 | 0 | 0 | 0 |
| FS2112 | $1.74 \times 10^{7}$ | $1.74 \times 10^{6}$ | 0 | 1 | 2 | 1 | $5.75 \times 10^{-7}$ |
| FS2130 ($RIF^{R}$ ARR-3) | $1.77 \times 10^{7}$ | $1.77 \times 10^{6}$ | L | L | L | L | 1 |
| FS2155 | $1.25 \times 10^{7}$ | $1.25 \times 10^{6}$ | 2 | 0 | 0 | 0.66666667 | $5.33 \times 10^{-7}$ |
| FS2258 | $1.42 \times 10^{7}$ | $1.42 \times 10^{6}$ | 0 | 0 | 0 | 0 | 0 |

**Supplementary Figure 1: Average nucleotide identities (ANI) and single nucleotide polymorphisms (SNPs) across all 13 paired isolates.**

ANI was calculated using the BLAST alignment algorithm (ANIb). The core genome SNPs are displayed in each cell below the ANI %, this was calculated using snippy v4.6.3 and snp-dist using FSFC1386 as reference.

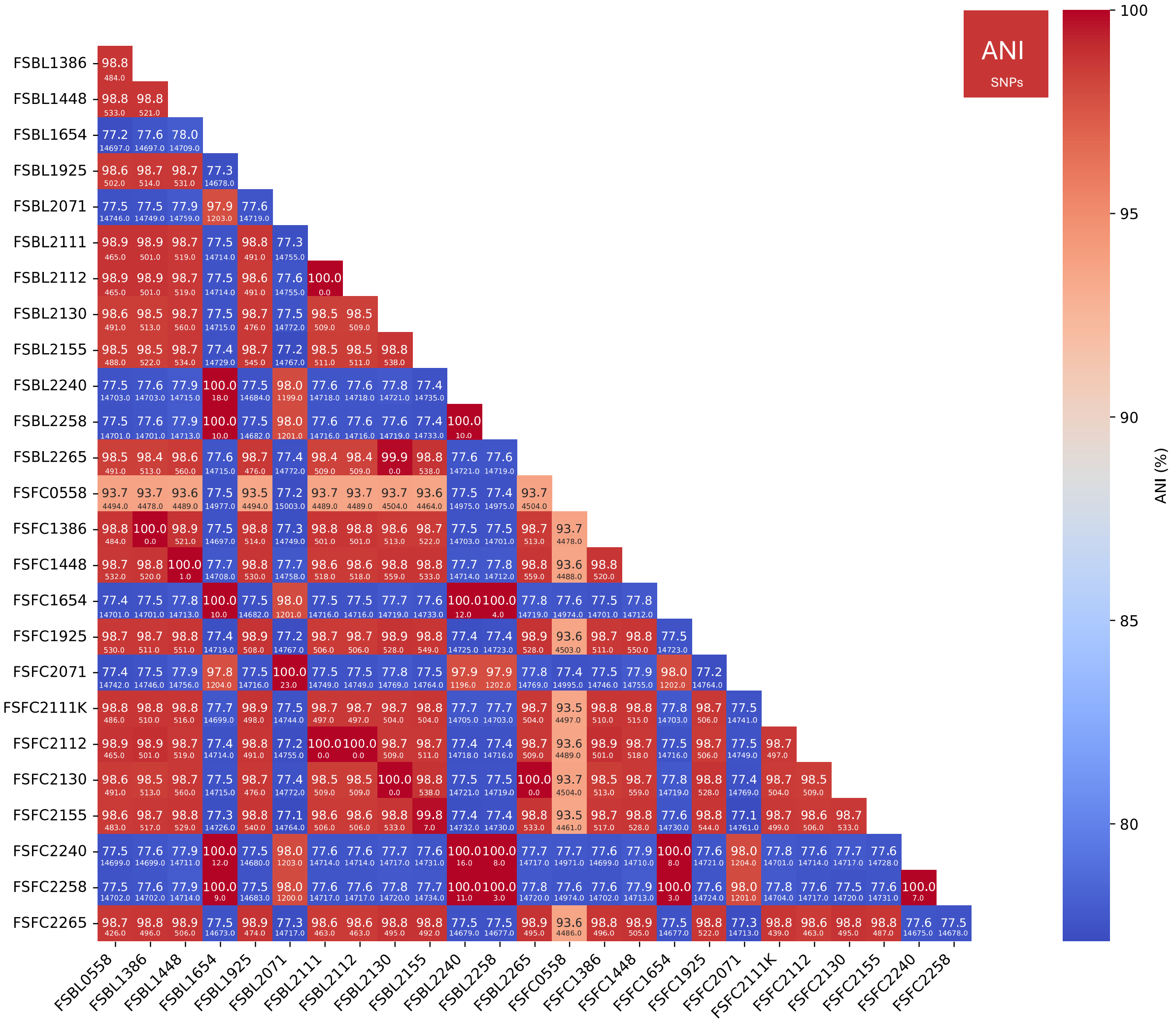

**Supplementary Figure 2: Loss and gain of genetic regions between contigs of highly related pairs**

Connections from Figure 4 were analysed and any contigs or regions of contigs not shared were visualised by snapgene. In FSFC1386 there is a 6415bp contig (FC plasmid 2) that is not in FSBL1386, this contains mainly phage genes. FS1448 and FS2071 contain contigs with phage related genes in the blood isolate but not in the faecal isolate. Plasmid 7 and 8 of the faecal isolate FSFC2130 are not found in the blood isolate FSBL2130, faecal plasmid 7 contains an IS*1* Tn, whereas faecal plasmid 8 contains only unknown genes. However, the blood isolate FSBL2130 contains a plasmid not found in the faecal isolate (BL p5) which harbours a *rep* protein and hypothetical proteins.

Over half (45.7%; 135,460bp/296,347bp) of the length of plasmids BL p1 and FC p1 of FS2071 do not align at 99.9% sequence identity, this means it is likely that there is loss of entire sections or large genomic movement such as recombination.

Between isolates of FS2258, blood plasmid 4 and faecal plasmid 4 do not align but this is due to a rearrangement in a replication protein.

The blood and faecal genomes of FS2155 have the largest disparities of all paired samples, with a 400kb disparity in chromosome size however this could be explained by the insertion of FC p1 into the chromosome. FSBL2155 p2 is mainly accounted for in FC p2, however the blood isolate has two extra plasmids BL p1 and BL p3. BL p1 is a 297kb plasmid containing IncF conjugative *tra* genes and the vap toxin producing gene. BL p1 shares the *fec* operon, encoding the Iron(III) dicitrate transport system, and the *lac* operon with FC p2. BL p3 is a 100kb plasmid with *tra* genes and a type I restriction-modification system, it shares an IS*26* with FC p2 seen as the connection on Figure 4 but no additional homology.

FS1654 and FS2240 were visualised differently based on the method of assembly for the faecal isolates (unicycler rather than hybracter) which is why the contigs are named different (e.g. c1 to c28 for FS1654), with some of these contigs being in the 100s of bp. With FS1654 there is one additional contig in the faecal c18 and 22 additional small contigs (p6 to p27) in the blood. However, this is likely a fragmented set of FC c18 as this contig aligns to these 15 small BL contigs at between 98.942% and 99.742%. FC c18 is 1552bp in length and contains a *rep* gene and hypothetical protein gene, the 15 small blood contigs contain both genes or just the *rep* gene. There are 7 contigs from the blood isolate FSBL1654 which share no homology with the faecal isolate FSFC1654, one of these is BL p7 which contains a *dfrA17* gene, previously shown to be the one difference in AMR profiles across the highly related pairs (see Supplementary Table 3), the rest are small contigs with either hypothetical genes or no genes.

With FS2240 there are additional contigs in the faecal isolates: c16 and c7. However, c7 aligns to the BL chromosome with 99.875% identity across its length (16,812bp) and c16 aligns to the chromosome with 99.647% across its length (851bp) likely meaning these were misassembled parts of the chromosome. FC c7 contains 24 genes, all phage or pro-phage related including those involved in the assembly of the phage head and tail and FC c16 contains two genes a phage antitermination protein gene and a Holliday junction resolvase gene: *rusA*. It is well known that phage genes being repetitive in nature are difficult to assemble if the reads don’t span the length of the longest repeat.

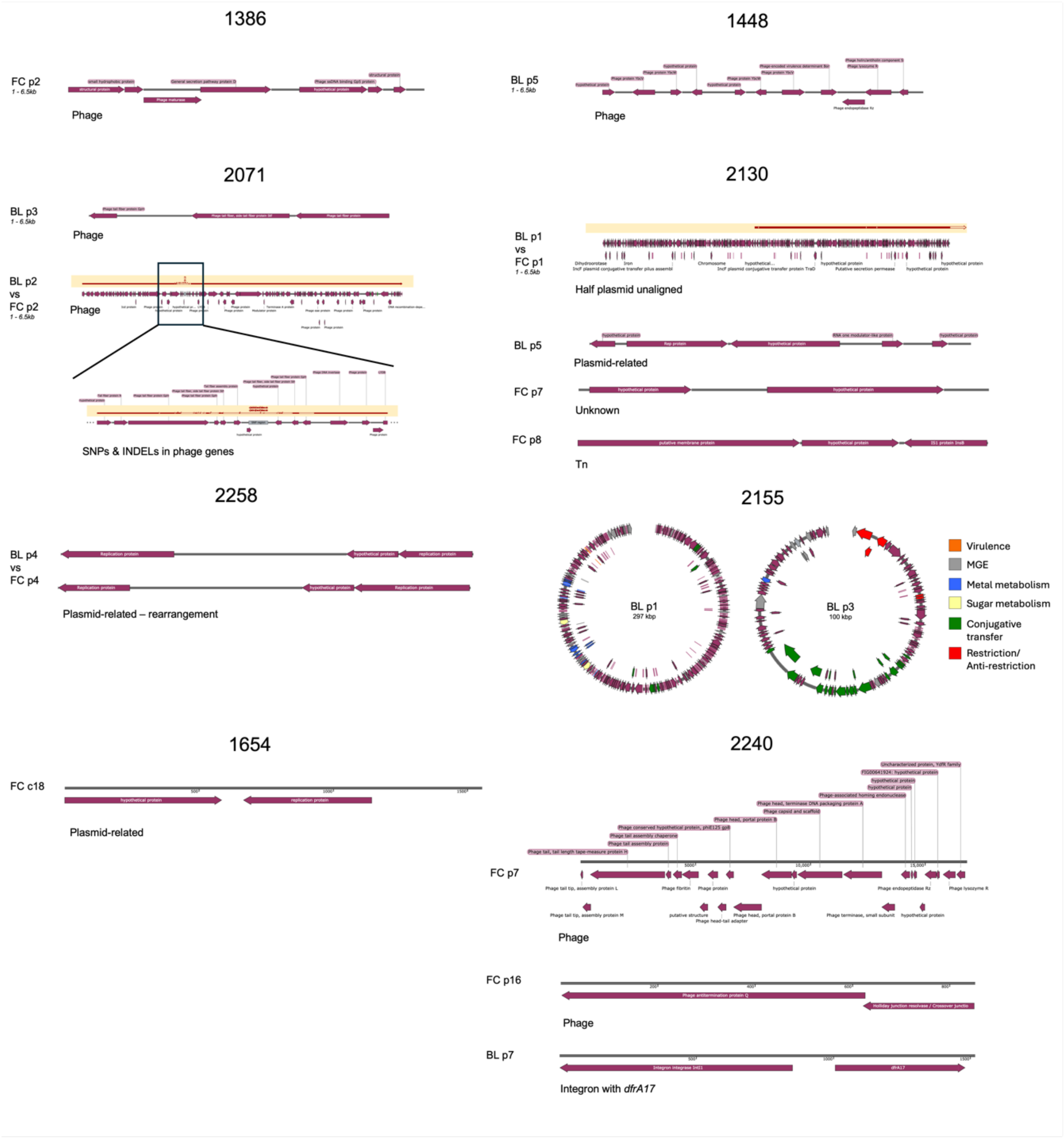
1386
FC p2
1 - 6.5kb
Phage
1448
BL p5
1 - 6.5kb
Phage
2071
BL p3
1 - 6.5kb
Phage
BL p2
vs
FC p2
1 - 6.5kb
Phage
SNPs & INDELs in phage genes
2130
BL p1
vs
FC p1
1 - 6.5kb
Half plasmid unaligned
BL p5
Plasmid-related
FC p7
Unknown
FC p8
Tn
2258
BL p4
vs
FC p4
Plasmid-related – rearrangement
2155
BL p1
297 kbp
BL p3
100 kbp
Virulence
MGE
Metal metabolism
Sugar metabolism
Conjugative transfer
Restriction/ Anti-restriction
1654
FC c18
Plasmid-related
2240
FC p7
Phage
FC p16
Phage
BL p7
Integron with *dfrA17*
